# High burden of premature ventricular contractions upregulates transcriptional markers of inflammation and promotes adverse cardiac remodeling linked to cardiomyopathy

**DOI:** 10.1101/2025.05.23.652709

**Authors:** J M L Medina-Contreras, Jaime Balderas-Villalobos, Jose Gomez-Arroyo, Janée Hayles, Karoly Kaszala, Alex Y Tan, Montserrat Samsó, Jose F Huizar, Jose M. Eltit

## Abstract

Premature ventricular contractions (PVCs) are the most prevalent ventricular arrhythmia in adults. High PVC burden can lead to left ventricular (LV) systolic dysfunction, eccentric hypertrophy, and an increased risk of heart failure (HF) and sudden cardiac death (SCD). Inadequate *angiogenesis* is a key determinant in the transition from adaptive to maladaptive cardiac hypertrophy and *fibrosis* is a risk factor for arrhythmia and SCD. To quantitatively assess structural remodeling and transcriptional alterations in PVC-induced cardiomyopathy (PVC-CM), animals were implanted with modified pacemakers to deliver bigeminal PVCs (200-220 ms coupling interval) for 12 weeks. Collagen deposition and interstitial ultrastructure of LV samples were analyzed using light and transmission electron microscopy, respectively. Pericytes, fibroblasts, myocytes, smooth muscle, and endothelial cells were imaged using confocal microscopy, quantified with an artificial intelligence-based segmentation analysis, and compared using hierarchical statistics. Transcriptional changes were assessed via RNAseq. Although cardiomyocytes hypertrophied in PVC-CM, capillary rarefaction was overcome by an increase in capillary-to-myocyte ratio. Additionally, thicker blood vessels were more abundant in PVC-CM. Fibroblast-to-myocyte ratio more than doubled, interstitial collagen fibers increased, and interstitial space thickened in PVC-CM. Transcripts involved in interstitial remodeling, inflammatory response, and alarmins were strongly elevated in PVC-CM. Overall, while the angiogenic response meets the metabolic demands of cardiac hypertrophy, upregulated markers of inflammation and cardiomyopathy linked to reactive fibrosis collectively represent an adverse LV remodeling that heightens the risk of HF and SCD in PVC-CM.

## Introduction

Premature ventricular contractions (PVCs) are the most prevalent ventricular arrhythmia in adults and often affect the elderly population^1, 2^. The sporadic occurrence of PVCs is believed to be benign, however frequent PVCs have been recognized as a cause of heart failure and left ventricular (LV) systolic dysfunction known as PVC-induced cardiomyopathy (PVC-CM)^2, 3^. Importantly, LV dysfunction together with high PVC burden have been associated with an increased risk of mortality, ventricular arrhythmias, and sudden cardiac death (SCD)^4–6^.

PVCs are ectopic ventricular beats that originate in ventricular tissue and result in dyssynchronous contraction due to a slow myocardial activation. High-burden PVCs progressively lead to cardiac remodeling characterized by decreased LV contractility (systolic dysfunction), electro-mechanical latency, and eventually, eccentric hypertrophy and LV dilatation^7, 8^. Whereas large animal models have been instrumental to systematically demonstrate that frequent PVCs cause PVC-CM^9, 10^, the mechanisms whereby frequent PVCs increase mortality are poorly understood^4, 5^.

Structurally, PVC-CM hearts show eccentric hypertrophy characterized by increased LV mass index with decreased relative wall thickness^8^. Additionally, frequent PVCs also induce interstitial (or diffuse) fibrosis in the ventricle^10–12^. LV hypertrophic responses are associated with neo-angiogenesis, likely as a compensatory mechanism to support the increased energy demand. Indeed, inadequate angiogenesis is a key step in the transition from an *adaptive* to *maladaptive* (or decompensated) cardiac hypertrophy^13–17^. Here, we quantitatively evaluate fibrotic and angiogenic responses, along with underlying transcriptional adaptations, in a translational PVC-CM model.

## Methods

### Animal model

This study conforms to the Guide for the Care and Use of Laboratory Animals approved by the Institution Animal Care and Use Committee at the Central Virginia HCS. Female mongrel canines (>10 months old, ∼21 Kg) were implanted with a modified dual-chamber epicardial pacemaker via left thoracotomy^9^. Surgeries were performed under general anesthesia using acepromazine (0.5-2 mg/kg, p.o.) and brevital (6-10 mg/kg, IV) for induction, and inhaled isoflurane (2-3%, endotracheal intubation) for maintenance throughout the procedure. Penicillin G 900,000 units IM were administered 12 hours prior to surgery. Postoperatively, buprenorphine (0.1-0.2mg/kg, IM) twice daily for 3 days was administered for pain control. After a 2-week surgical recovery, animals were randomized into 2 groups: 1) PVC-CM group, receiving bigeminal PVCs (50% burden) with a 200-220 ms coupling interval for 12 weeks, and 2) sham group where PVCs were not enabled. Echocardiographic recordings were obtained at baseline and at 12-weeks of chronic PVCs to evaluate the development of cardiomyopathy. The tissue samples used in this study were obtained from the same experimental animal groups described in our previous publication^18^. Euthanasia was performed at week 12, via exsanguination during final left thoracotomy surgery under general anesthesia. The harvested heart was immediately rinsed with ice-cold phosphate-buffered saline (PBS), which was injected directly through the left main coronary artery to remove blood. Segments of the anterior LV were: 1) snap frozen in liquid nitrogen for biochemical analysis, 2) fixed in 10% neutral buffered formalin (NBF) for optical microscopy, and 3) perfused first with 2 mM EGTA and 16 mM KCl in PBS, followed by fixation in 4% glutaraldehyde, 0.1M cacodylate, pH 7.4 for transmission electron microscopy. Freshly fixed tissue samples were utilized in immunostaining experiments to prevent damage caused by freezing. However, due to the limited availability of fresh samples as the project progressed, the number of animals used for different molecular markers varied.

### RNA sequencing of LV samples

Snap-frozen myocardial tissue from randomly selected regions of the left ventricular (LV) free wall was used for RNA extraction. Total RNA was isolated using the RNeasy kit (Qiagen, MD, USA) to generate fourteen ribo-depleted, paired-end, 75 base-pair stranded RNA libraries (6 samples from sham animals and 8 from PVC-CM animals). Samples were sequenced using an Illumina NextSeq sequencing platform. Reads were pseudo- aligned and quantified using an index transcriptome version of the CanFam3.1 dog genome assembly (GCA_000002285.2) using Kallisto with standard settings^19^. Transcript-level abundance estimates were imported and summarized as a counts matrix using tximport^20^. Gene-level exploratory analysis and differential gene expression was performed using DESeq-2 R with standard settings, setting a Benjamini and Hochberg false discovery rate of <0.1^21^. Data have been deposited in the Gene Expression Omnibus (GEO) public repository under accession number GSE296225.

### Statistical Analysis

Data distribution of Western blots was determined using the Kolmogorov–Smirnov test. Unpaired *t*-test or Mann–Whitney test was used to compare differences between groups for parametric or nonparametric distributions, respectively. When multiple measurements were collected per animal (AIVIA analysis and in situ Sirius Red / Fast green collagen determination) nested *t*-test, a hierarchical statistical method, was used for comparison of the data^22, 23^. The number of animals (N) studied in each experiment is indicated in the corresponding figure or in Table. In addition, for hierarchical statistical analyses, data were represented using box and whiskers plot along with individual values. Hierarchical statistics (nested t-test) account variability within and between subjects^22, 23^. The threshold for the significant difference was *p* < 0.05. GraphPad Prism 9.3.1 was used as a statistical software resource.

Additional methods are described in supplementary material.

## Results

### Cardiomyocyte hypertrophy

Previously, we reported the development of eccentric hypertrophy^8^ and an increase in cardiomyocyte size^18^. The cohorts of animals used in the latter work are the same as those used here, where the echocardiographic results showed 44.36 ± 5.31% vs 61.91 ± 5.61%**** LV ejection fraction (LVEF), in the 12-week-PVC-CM and sham cohort, respectively (Šídák’s multiple comparisons test \*\*\*\**p* < .0001)^18^. Our extended determinations and analysis of in situ cross-sectional myocyte size (Table 1) refine previous measurements and show that the cross-sectional area of myocytes in the PVC- CM group is 38% larger than in shams. Considering the average mean value per micrograph, the number of animals studied, and the number of myocytes measured (20,287 and 21,070 myocytes in the PVC-CM and sham groups, respectively; see Table 1), this difference is statistically significant using hierarchical statistics (p<0.001, nested t-test).

**Table 1.**
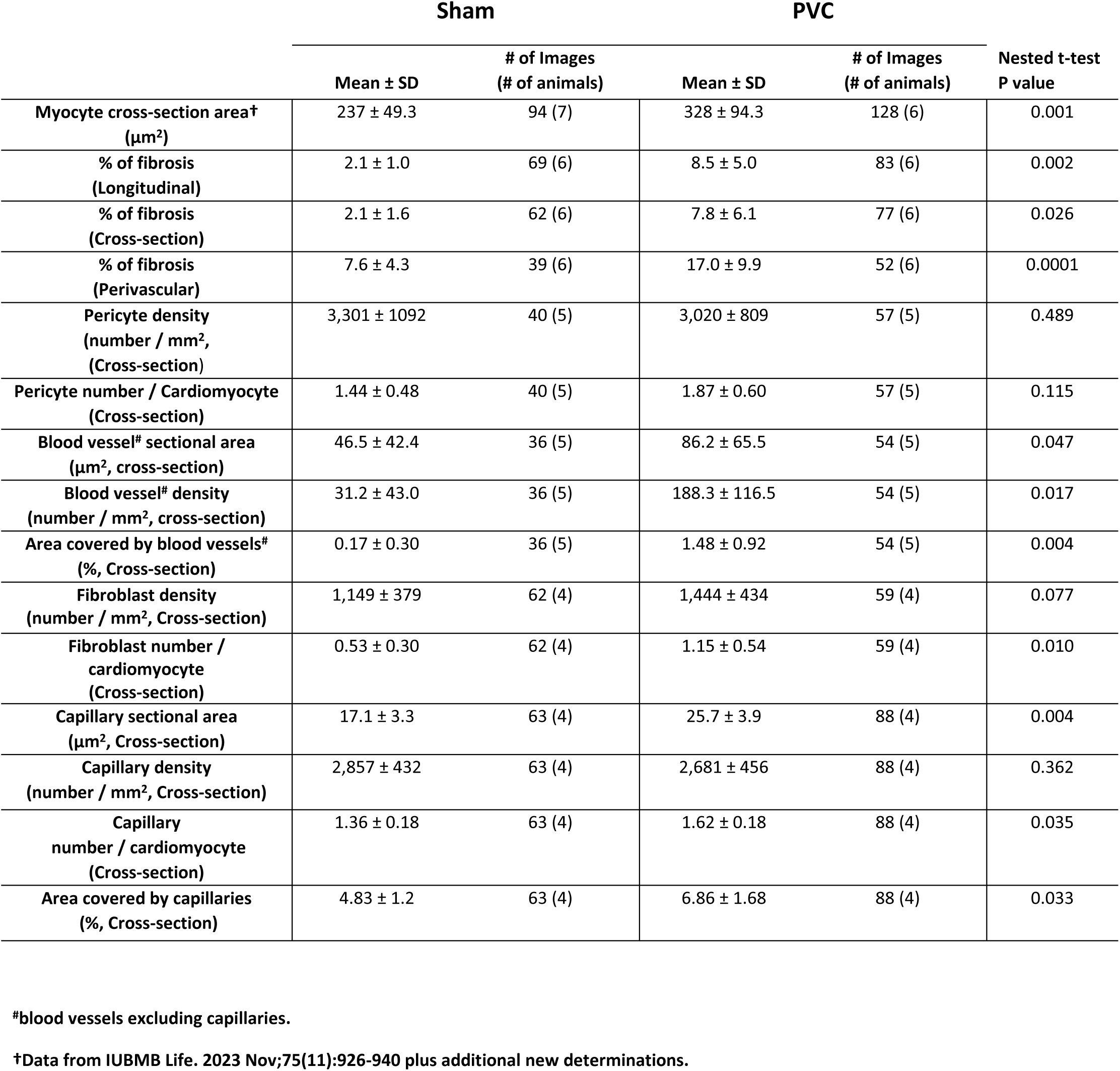
Quantification of cellular composition of LV tissue.

### Interstitial and perivascular fibrosis

Fibrosis was assessed by staining collagen in situ and followed by visualization under light microscopy. The PVC-CM group showed an increased percentage of collagen deposition compared to the sham group (Fig. 1A). Specifically, in longitudinal sections, the PVC-CM group showed a 4-fold increase in fibrosis compared to the sham group. In cross-sections the PVC-CM group showed a 3.8-fold increase, while perivascular fibrosis increased by 2.2-fold (Fig. 1A and Table 1). Cross-sections were also studied under the electron microscope to visualize ultrastructural changes of the interstitial space. Cardiomyocytes and capillaries (endothelial cells) were clearly identified in the TEM images of fixed LV samples (Fig. 1B). Representative images in Figure 1B depict the entire interstitial space (highlighted in yellow), comprised by the narrow spaces between adjacent cardiomyocytes, and between cardiomyocytes and capillaries. The interstitial area increased in the PVC-CM group with respect to the sham, consistent with the increased interstitial fibrosis observed in PVC-CM using optical microscopy (Fig. 1A).

**Figure 1.**
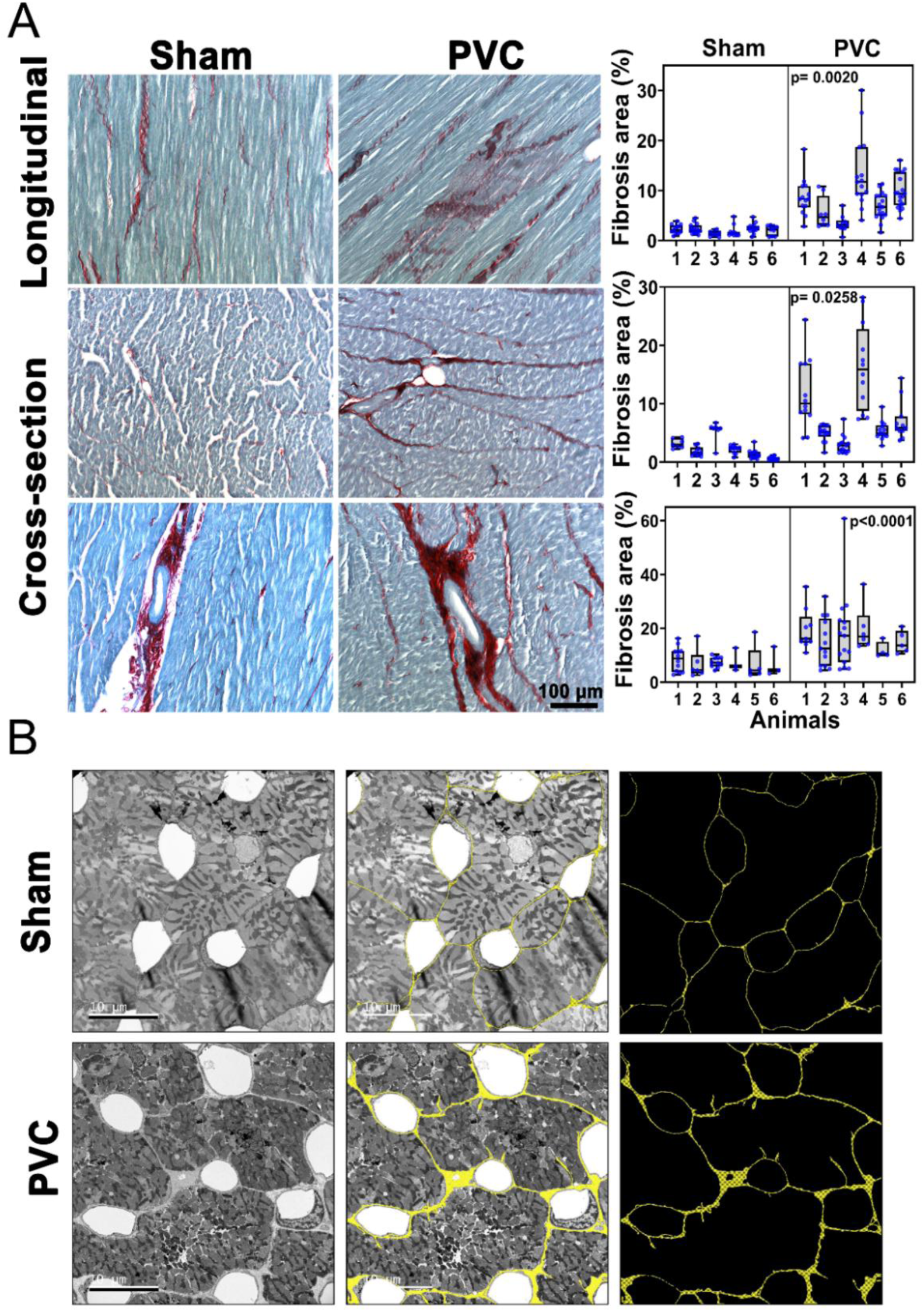
Fibrosis is elevated in PVC-CM. (A) Representative images of left ventricular (LV) free wall samples stained with Sirius Red / Fast Green to detect fibrotic depositions. Images were obtained from longitudinal (upper panels) and cross-sections (middle and lower panels) from sham and PVC-CM animals. The red staining reveals interstitial fibrosis (upper and middle panels) and fibrosis in perivascular regions (lower panels). Percentual area covered by collagen deposits was quantified; data from each animal are represented using box and whisker plots and data from each image are represented by blue dots (right graphs and Table 1). Both groups are compared using hierarchical statistics (nested t-test) and p values are shown in each graph. (B) Ultrastructural analysis using transmission electron microscopy (TEM). Representative TEM micrographs of LV thin cross-sections from sham and PVC-CM groups (left panels). The AIVIA software was used to highlight the interstitial space shown in yellow (merged in the middle panels); for comparison purposes the interstitial space is shown alone (right panels).

### Identification of cell types and vascular structures in LV tissue

To better characterize the cellular adaptations occurring in PVC-CM, a set of four markers was used to identify different cell types in the myocardium: α-smooth muscle actin (αSMA), vimentin (VIM), isolectin B4 (IB4), and DAPI. αSMA immunostaining unveiled abundant *pericytes*, thin and long cells in the interstitial space of the myocardium, depicted in red in the longitudinal sections of Figure 2A. In addition, as expected, the anti-αSMA antibody recognized *smooth muscle cells* forming the tunica media of blood vessels (see cross-section in Fig. 2A). Conversely, IB4 showed a selective stain for *endothelial cells* (green cells in Fig. 2A), which constitute both the capillaries and the tunica intima of larger blood vessels (Fig. 2A). The overlay of these markers revealed a delicate structural arrangement between the thin αSMA^+^ pericytes, which were intimately intertwined with IB4^+^ capillary endothelial cells, as observed in the merged image of the longitudinal section (Fig. 2A). This interaction resembles that of *pericytes* and *capillary endothelial cells* previously described in brain and heart^24–26^. Under our experimental conditions, the antibody targeting VIM, a recognized fibroblast marker ^27, 28^, strongly stained cells that were distinct from both αSMA⁺ pericytes and IB4⁺ capillary endothelial cells (white in Fig. 2A), indicating that VIM selectively identifies fibroblasts among interstitial cells. The VIM^+^ fibroblasts in the interstitial space contact both αSMA^+^ pericytes and IB4^+^ endothelial cells (Fig. 2A). In addition, the VIM antibody labeled endothelial cells in the tunica intima of large blood vessels, revealing a distinctive ring-shaped pattern within the inner layer (Fig. 2A cross-section), but was undetectable in endothelial cells forming capillaries. Overall, each selected cellular marker effectively distinguished the cell types of interest (Fig. 2A), confirming their suitability for quantitative analysis. Then samples from PVC-CM and sham groups were evaluated using these markers (αSMA, IB4, VIM, and DAPI) in cross- and longitudinal sections. Confocal images showed a clear increase in blood vessels in PVC-CM vs. sham for both cross- and longitudinal sections (Fig. 2B). Additionally, the wide extent and very well-organized interaction between αSMA^+^ pericytes and IB4^+^ capillary endothelial cells was clearly observed in longitudinal sections in both groups (Fig. 2B merge).

**Figure 2.**
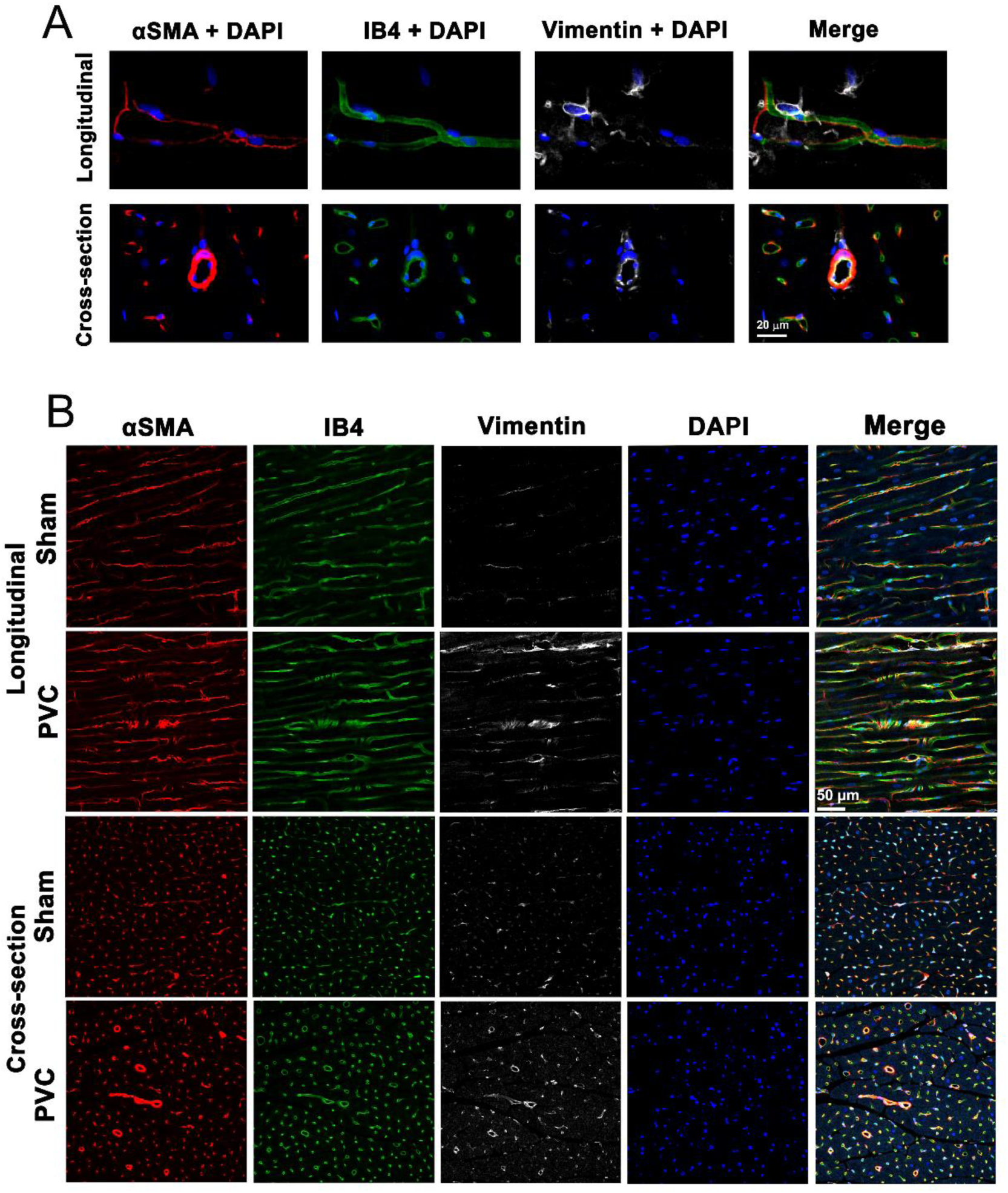
Identification of cells and structures in cardiac tissue using different cellular type markers. (A) Representative images showing formalin-fixed LV free wall tissue samples co-stained using anti-αSMA (red), anti-vimentin (VIM, white), isolectin B4 (IB4, green), and DAPI as counterstain (blue). In our experimental conditions, each marker recognized distinctive cell types and structures that were then quantified using the AIVIA software. In the longitudinal image (upper panels) αSMA^+^ cells show structure compatible with pericytes, IB4^+^ cells are structurally compatible with capillary endothelial cells, and VIM^+^ cells look like fibroblasts. Cross-section (lower panels) shows that anti- αSMA antibody stains the tunica media of blood vessels; IB4 stains capillaries and the tunica intima of blood vessels, and the anti-vimentin antibody stains the tunica intima of blood vessels and fibroblasts. (B) Representative images showing formalin-fixed LV free wall tissue samples of the two experimental groups co-stained with anti-αSMA, isolectin B4 (IB4), anti-vimentin (VIM) and DAPI as counterstain.

Myofibroblasts (as fibroblasts) are involved in collagen deposition but, in contrast to fibroblasts, are particularly important in scar compaction during replacement fibrosis ^29^. It is well known that fibroblast to myofibroblast differentiation requires an increase in expression of the contractile protein αSMA ^30, 31^. Myofibroblast presence, thus, may be captured by the colocalization of αSMA and VIM staining^32^. The αSMA^+^ and VIM^+^ cells present in the interstitium produced distinctive staining patterns with apparent low colocalization (Fig. 3, Table 2). Because αSMA and VIM can also colocalize in the boundary between the tunica media and the intima of blood vessels, the colocalization parameters between αSMA and VIM staining were determined ‘including’ and ‘omitting’ blood vessels. Persson’s and Mander’s correlation coefficients^33^ did not show significant statistical differences between the sham and PVC-CM groups. The percentage of pixel overlap (regardless of their intensity) between channels was low in both groups and showed a ∼3-fold increase from 0.31% to 0.90% in PVC-CM vs. sham group, respectively (p=0.065 nested t-test, Table 2). Such an increase was nullified when blood vessels were omitted from the analysis, suggesting that the increase in pixel overlap is accounted for by an increase in blood vessels in the PVC-CM group (Table 1). This analysis suggests that αSMA is not overexpressed in VIM^+^ fibroblasts, further suggesting limited trans differentiation toward a myofibroblast phenotype in the PVC-CM group.

**Figure 3.**
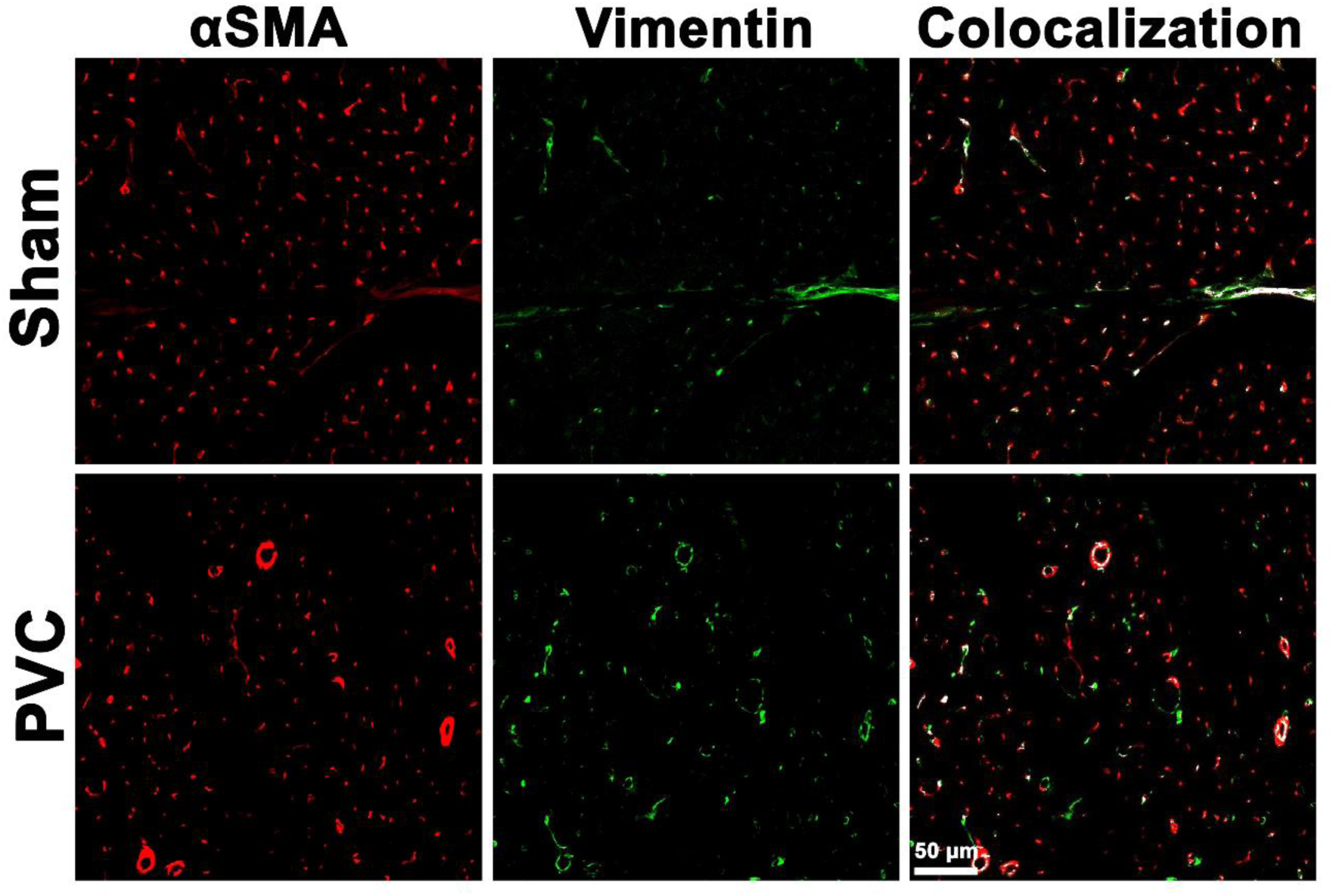
Low level of colocalization between αSMA and vimentin imply low presence of myofibroblasts. Fibroblasts, identified by vimentin-positive staining, gain αSMA expression during their differentiation to myofibroblasts. Representative images showing cross-sections of LV free wall tissue samples co-stained using anti-αSMA (red) and anti-vimentin (green). Overlaid pixels are represented in white. Co-localization parameters were further calculated using *Colocalization Finder* plug-in of the *Image J* software. As blood vessels react to both stains, colocalization parameters with and without including structures compatible with blood vessels were computed (see Table 2).

**Table 2.** Colocalization parameters for vimentin and α-smooth muscle actin in co- stained tissue.

| Whole tissue | # Animals | # Images | Pearson's R | M1* norm | M2* norm | %pixels |
| --- | --- | --- | --- | --- | --- | --- |
| Sham | 3 | 41 | $-0.082 \pm 0.144$ | $0.367 \pm 0.146$ | $0.122 \pm 0.064$ | $0.31 \pm 0.26$ |
| PVC | 3 | 71 | $-0.104 \pm 0.089$ | $0.484 \pm 0.103$ | $0.212 \pm 0.098$ | $0.90 \pm 0.55$ |
| nested t-test |  |  | p = 0.646 | p = 0.257 | p = 0.204 | p = 0.065 |

| Removing Blood Vessels | # Animals | # Images | Pearson's R | M1* norm | M2* norm | %pixels |
| --- | --- | --- | --- | --- | --- | --- |
| Sham | 3 | 41 | $-0.140 \pm 0.080$ | $0.294 \pm 0.103$ | $0.069 \pm 0.053$ | $0.10 \pm 0.07$ |
| PVC | 3 | 71 | $-0.158 \pm 0.085$ | $0.311 \pm 0.084$ | $0.096 \pm 0.054$ | $0.18 \pm 0.10$ |
| nested t-test |  |  | p = 0.424 | p = 0.816 | p = 0.427 | p = 1.052 |

\* Mander's correlation coefficients.

### Morphometric assessment of pericytes and vasculature

As shown in Figure 2A, αSMA was expressed mainly in pericytes and vascular smooth muscle cells with apparent lack of expression in VIM^+^ fibroblasts (Fig. 3). First, Western blot was used to quantify αSMA in whole LV tissue lysates, the PVC-CM group showed a 64% increase in expression compared to the sham group (p<0.0001 t-test, n=6 animals per group, Fig. 4A). Western blot shows the total expression of the protein, regardless of the cell type in which it is expressed. Then, to better estimate the subpopulation of cells responsible for the increase in αSMA, we performed quantitative morphometric determinations of confocal images stained with anti-SMA antibody using a segmentation function in the AIVIA software. For αSMA staining in cross sections, the pattern was split into two: the αSMA^+^ pericytes and the distinctive ring-shaped tunica media (smooth muscle layer) of blood vessels (Fig. 4B). In the same confocal image, the cell contour was stained using wheat germ agglutinin (WGA) and the AIVIA software was trained to identify the total number of myocytes (Fig. 4B). This procedure allowed us to quantify the number of pericytes, the number of blood vessels (excluding capillaries), and the total number of myocytes per image. This quantification was applied to several images per animal in the PVC-CM and sham groups (Fig. 4C). The density of pericytes (pericyte/mm^2^) was similar in PVC-CM and the sham group (Table 1 and Fig. 4C, p=0.489, nested t-test). After accounting for the hypertrophy-induced increase in myocyte size in the PVC-CM group^18^ (see Table 1), the calculated pericyte-to-cardiomyocyte ratio revealed a trend of approximately 30% more pericytes per myocyte in the PVC-CM group, that was not statistically significant (Table 1 and Fig. 4C, p=0.115, nested t-test). The PVC-CM group showed blood vessels (excluding capillaries) with 86% larger sectional area than the sham group (Table 1 and Fig. 4C, p=0.047 nested t-test). In addition, the cross-sectional images revealed that the density of blood vessels (number per mm^2^) and the percentual area covered by them were 6-fold (p=0.017, nested t-test) and 8.7-fold (p=0.004, nested t-test) larger, respectively, in PVC-CM when compared to the sham group (Table 1 and Fig. 4C).

**Figure 4.**
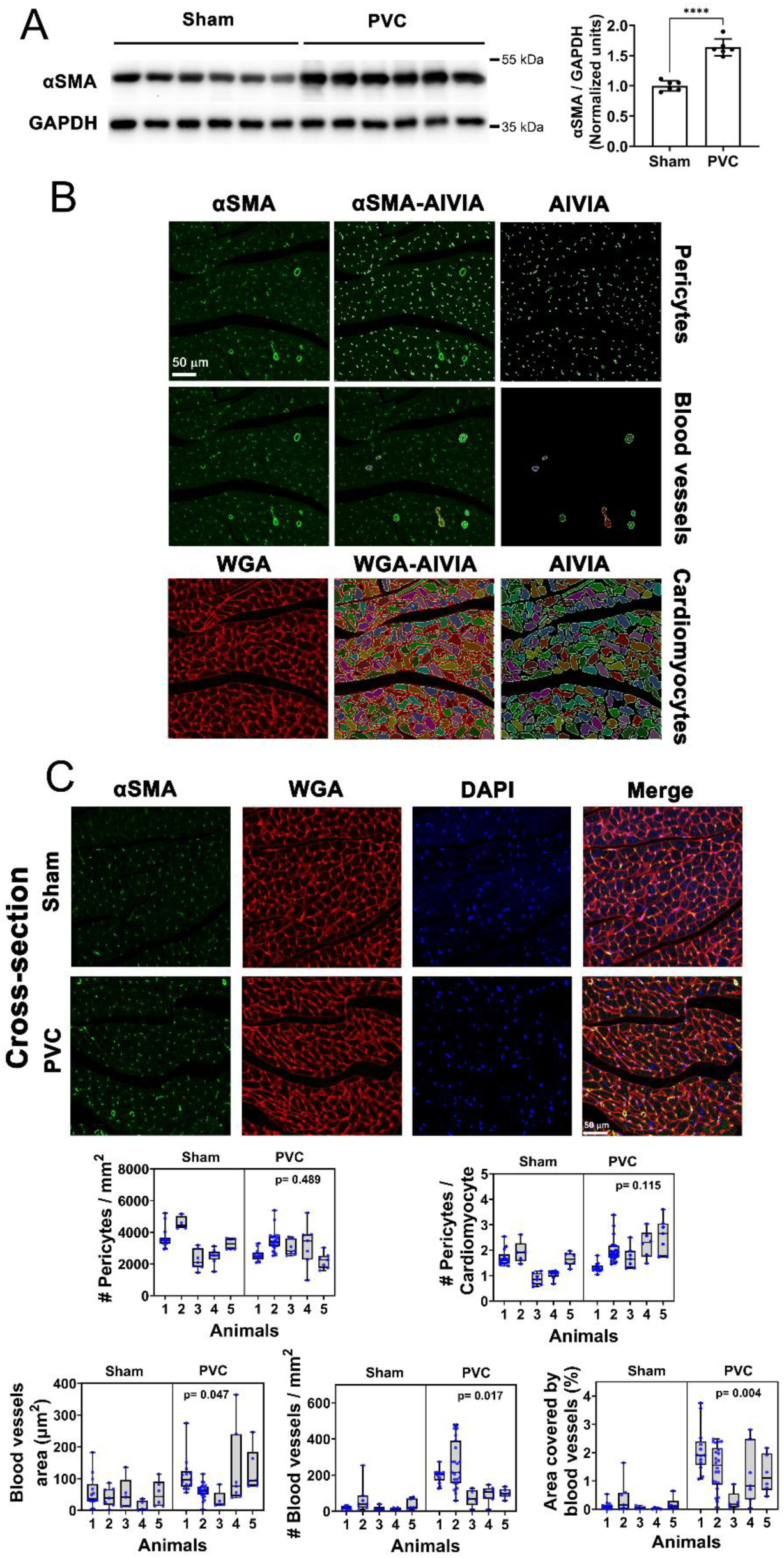
Protein expression and morphometric analysis of αSMA revealed pericytes and blood vessels in LV cardiac tissue. (A) Western blot analysis shows increase in αSMA expression in PVC-CM vs. sham (p<0.0001 t-test, n=6 animal per group). (B) Representative LV cross-sections co-stained with anti-αSMA antibody (green) and WGA-AF633 (red) (left panels). Artificial intelligence-based (AIVIA) recognition and segmentation of αSMA^+^ cells (pericytes) and blood vessels, and WGA-stained cardiomyocytes overlayed with the original image (middle panels). AIVIA generated regions of interest (ROIs) corresponding to pericytes, blood vessels, and cardiomyocytes (right panels). (C) Representative formalin-fixed LV cross-sections co-stained using anti- αSMA (green), WGA-AF633 (red), and DAPI (blue) in sham and PVC-CM LV free wall samples. The identified ROIs were used to calculate the number of pericytes, number and area of blood vessels, and number of myocyte (WGA) on each image. AIVIA results are plotted and analyzed using hierarchical statistics (nested-t test). The density (number of pericytes per mm^2^) and number of pericytes per cardiomyocyte were quantified for several images per animal. In total 13,529 vs. 17,641 pericytes were analyzed for sham vs. PVC, respectively. The unitary area of blood vessels (μm^2^), density (number of blood vessels per mm^2^) and the percentage of area covered by blood vessels were also quantified using the artificial intelligence-based software. The total number of blood vessels analyzed was 36 vs. 54 in the sham vs. PVC-CM group, respectively. Several micrographs were quantified per animal (see Table 1). Each point in the graphs represents the mean value per micrograph, and the data are displayed as a box and whisker plot.

### Morphometric analysis of fibroblasts

VIM, that was shown to be expressed in fibroblasts and the intima of blood vessels (Fig 2A), was strongly elevated when assessed using Western blot in 4 out of 6 PVC-CM samples, showing collectively a 139% increase compared to the sham group. However, this trend was not statistically significant, p=0.085 t-test, due to high data dispersion (Fig. 5A). Besides, interleukin 1β (IL-1β), an inflammatory cytokine associated with cardiac remodeling and fibrosis^34, 35^, showed a significant 69% increase in expression in PVC- CM samples compared to the sham group (p=0.007, t-test, Fig. 5A).

**Figure 5.**
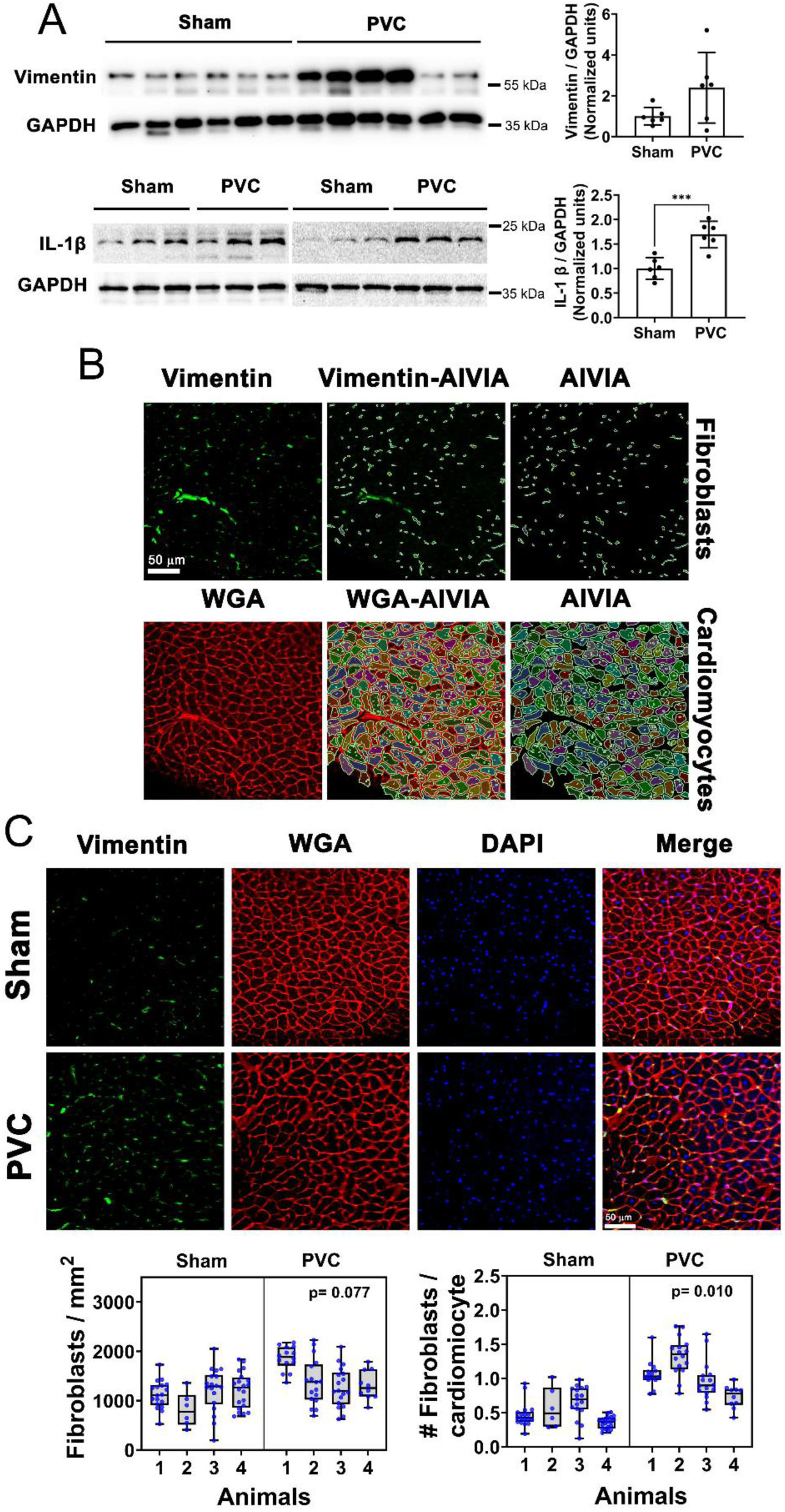
Frequent PVCs increase fibroblast content. (A) Western blot study was used to measure vimentin expression in the LV tissue in PVC-CM with respect to sham (p=0.085 t-test, n= 6 animals per condition) and IL-1β (p=0.007 t-test, n= 6 animals per condition). (B) Original representative LV cross-sections co-stained with anti-vimentin antibody (green) and WGA-AF633 (red) (left panels). AI-powered image segmentation merged with the representative image (middle panels). The segmentation result showed fibroblasts and cardiomyocytes, while blood vessel structures were rejected (right panels). (C) Representative LV cross-sections co-stained using anti-vimentin (green), WGA-AF633 (red), and DAPI (blue) in sham and PVC-CM samples. The ROIs identified were used to calculate the number of fibroblasts and the number of myocytes on each image. Results from the AIVIA software were plotted and analyzed using hierarchical statistics (nested t-test). Density (number of fibroblasts per mm^2^) and number of fibroblasts per cardiomyocyte were quantified for several images. The number of fibroblasts analyzed was 7,305 and 7,407 for sham and PVC-CM group, respectively. Each point in the graphs represents the mean value per micrograph, and the data is displayed as a box and whisker plot.

The AIVIA software was trained to identify the staining pattern consistent with VIM^+^ fibroblasts (i.e., excluding the blood vessel staining pattern) and the total number of myocytes in the corresponding WGA channel from the same confocal image (Fig. 5B). The density of fibroblasts (fibroblasts/mm^2^) in the images was 26% larger in PVC-CM samples than in sham, which almost reached statistical significance (Table 1 and Fig. 5C, p=0.077, nested t-test). Importantly, the PVC-CM group doubled (2.2-fold) the number of fibroblasts compared to the sham group when adjusted per number of myocytes (Table 1 and Fig. 5C, p=0.010, nested t-test), suggesting the active proliferation of fibroblasts in PVC-CM.

### Morphometric analysis of capillaries

We investigated the potential occurrence of capillary rarefaction, which is a key characteristic of maladaptive hypertrophy. First, western blot analysis of whole tissue samples was carried out, with special emphasis on proteins involved in the function and remodeling of capillaries. Both, the endothelial nitric oxide synthase (eNOS), a key protein involved in the generation of NO that controls smooth muscle relaxation expressed in endothelial cells^36^, and the vascular endothelial growth factor-B (VEGF-B), a strong angiogenic growth factor^37^, are elevated in the PVC-CM LV samples compared to the sham group (21% increase, p<0.036, and 34% increase p<0.003, respectively vs. sham, t-test) (Fig. 6A). Then, IB4 staining was used to identify endothelial cells (Fig. 2A). The AIVIA software was trained to identify endothelial cells forming capillaries but reject the ones forming the intima of blood vessels (Fig. 6B). Additionally, WGA staining was used to compute the total number of myocytes in the same confocal image (Fig. 6B). Interestingly, the caliber of the capillaries is larger in the PVC-CM samples than in the sham group (Fig. 6C). Morphometric assessment showed that the PVC-CM group had capillaries with 50% larger cross-sectional area than the sham group (Table 1 and Fig. 6C, p= 0.004 nested t-test). The density of capillaries (number of capillaries / mm^2^) was not different between the two experimental groups (Table 1 and Fig. 6C, p =0.362, nested t-test), but the PVC-CM group had 19% more capillaries per myocyte than the sham group (Table 1 and Fig. 6C, p= 0.035 nested t-test). In addition, the area covered by capillaries in the cross-section images was 42% larger in PVC-CM vs. the sham group (Table 1 and Fig. 6C, p= 0.033 nested t-test).

**Figure 6.**
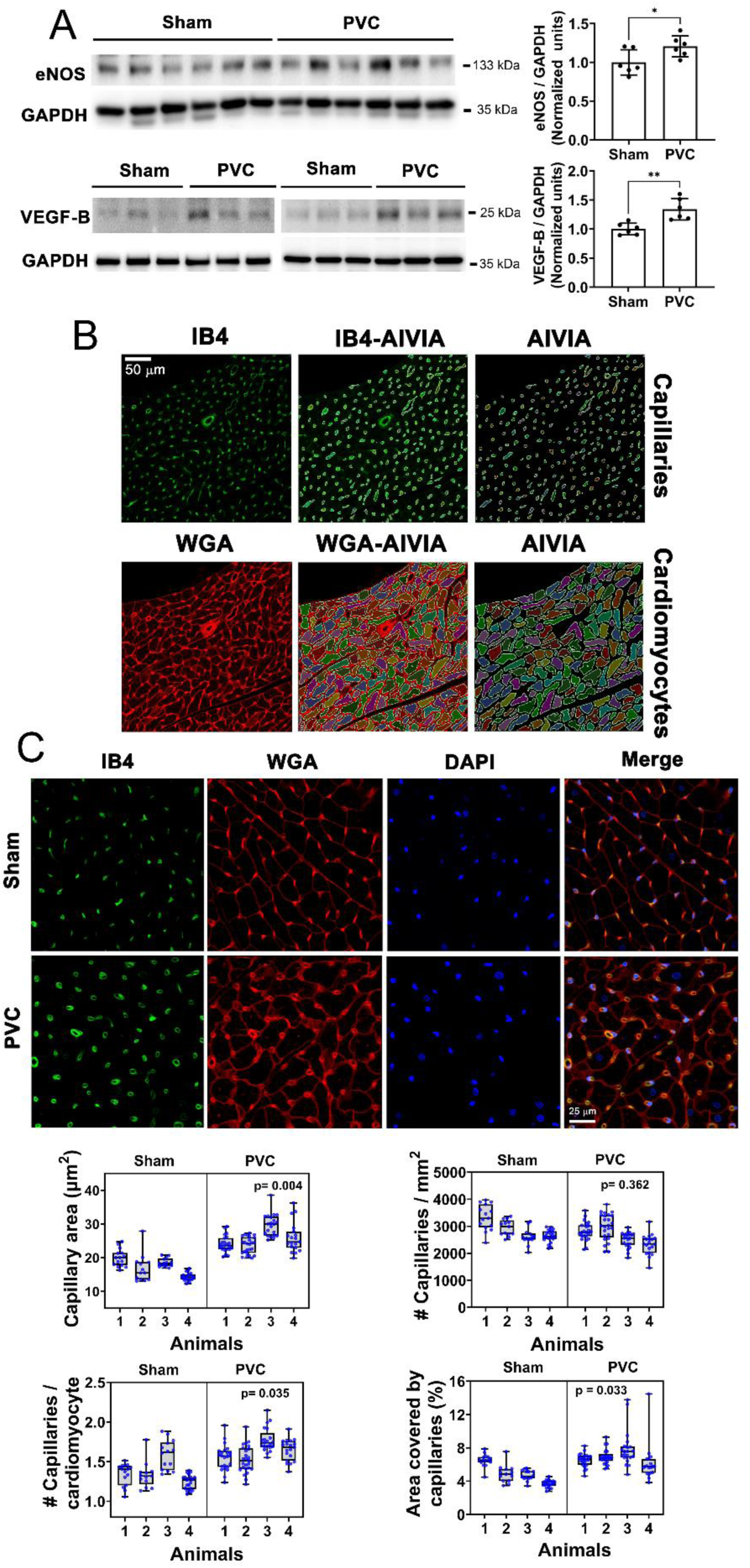
Frequent PVCs promote an angiogenic response. (A) Western blot analysis indicates increase expression of eNOS (p<0.036 t-test, n= 6 animals per condition) and VEGF-B (p<0.003 t-test, n= 6 animals per condition). (B) Original representative LV cross- sections co-stained with IB4-AF488 (green) and WGA-AF633 (red; left panels). AI-based detection of capillaries and cardiomyocytes using the AI-powered AIVIA classifier merged with the original images (middle panels). Capillaries and cardiomyocytes were recognized by the AI segmentation, while blood vessel structures were rejected (right panels). (C) Representative LV cross-sections co-stained with IB4 (green), WGA-AF633 (red), and DAPI (blue) in sham and PVC-CM. The recognized ROIs were used to calculate the number of capillaries, area of capillaries, and number of myocytes on each image. Results from AIVIA software were plotted and analyzed using hierarchical statistics (nested t-test) (Table 1). The unitary capillary area (μm^2^), density (number of capillaries per mm^2^), number of capillaries per cardiomyocyte, and the percentual area covered by capillaries were quantified for several images per animal. The number of capillaries analyzed was 18,318 vs. 24,012 in the sham vs. PVC-CM group, respectively. Several micrographs were analyzed per animal (data shown in Table 1) and the comparison between groups was performed using hierarchical statistical analysis (nested t-test). Each point in the graphs represents the mean value per micrograph, and the data are displayed as a box and whisker plot.

To better visualize the interaction between pericytes and capillaries, Figure 7A shows a TEM image from a PVC-CM LV sample depicting several contiguous capillaries with different sizes and interstitial cells (pericytes) interconnecting capillaries (arrows in Fig. 7A). In confocal cross-section images of the PVC-CM sample (Fig. 7B), several examples of αSMA^+^ cells (pericytes, displayed in red) were interacting either with single capillaries (in green) or connecting pairs of capillaries showing similar structural arrangement to that observed in the TEM image.

**Figure 7.**
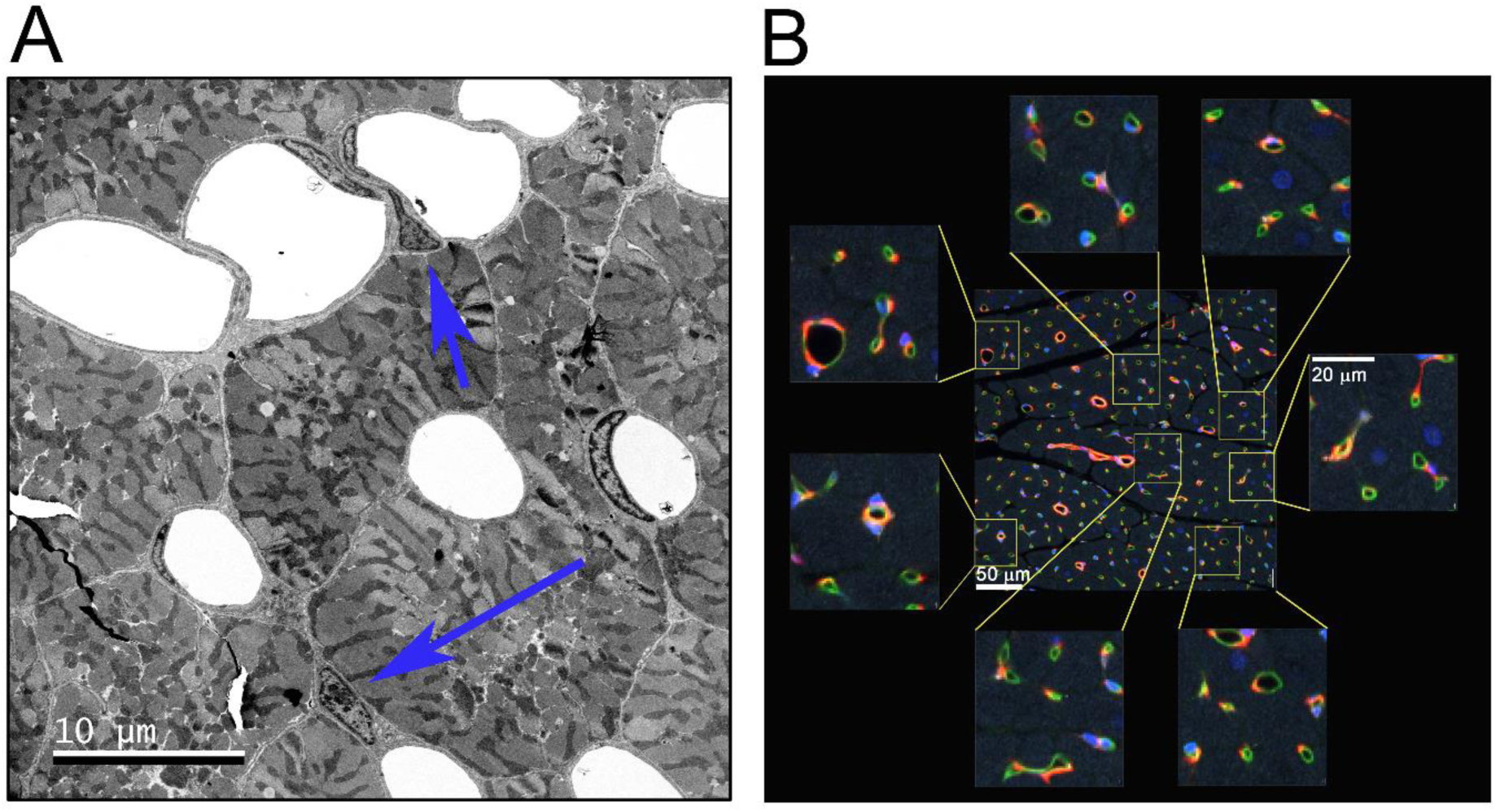
TEM and confocal microscopy reveal PVC-CM- induced alterations in capillary and pericyte organization. (A) Representative TEM micrograph of a PVC-CM sample showing contiguous capillaries with different caliber (arrangement rarely seen in the sham condition; see Fig. 1B for comparison). The blue arrows indicate the presence of pericytes. (B) Representative LV cross-section of a PVC-CM sample co-stained with IB4 (green; endothelial cells in capillaries and intima layer blood vessels), anti-αSMA (red; pericytes and tunica media of blood vessels), anti-vimentin (white, fibroblasts), and DAPI (blue, nuclei); image taken from Fig. 2B (merge). The enlarged quadrants show adjacent capillaries with some of them inter-connected by pericytes similar to these in the TEM image.

### RNA seq analysis of LV samples

Bulk RNA sequencing analysis was performed to investigate transcriptional changes and uncover potential mechanisms driving the cardiac adaptations observed in PVC-CM. A total of 646 (2.9%) and 470 (2.1%) genes were up- or down-regulated, respectively, in PVC-CM tissue compared to sham. The heatmap displaying the top 50 upregulated genes is shown in Figure 8A. Gene set enrichment analysis of the top 50 upregulated genes revealed significant associations with biological processes related to proteolysis, neutrophil chemotaxis, immune response, and extracellular matrix organization (Fig. 8B). The transcriptional signature also showed enrichment for transcriptional regulatory networks associated with nuclear factor kappa-light-chain-enhancer of activated B cells (NF-κB) or RELA (which encodes a subunit of NF- κB), as determined by transcriptional regulatory relationship analysis (using TRRUST5 software; Fig. 8C).

**Figure 8.**
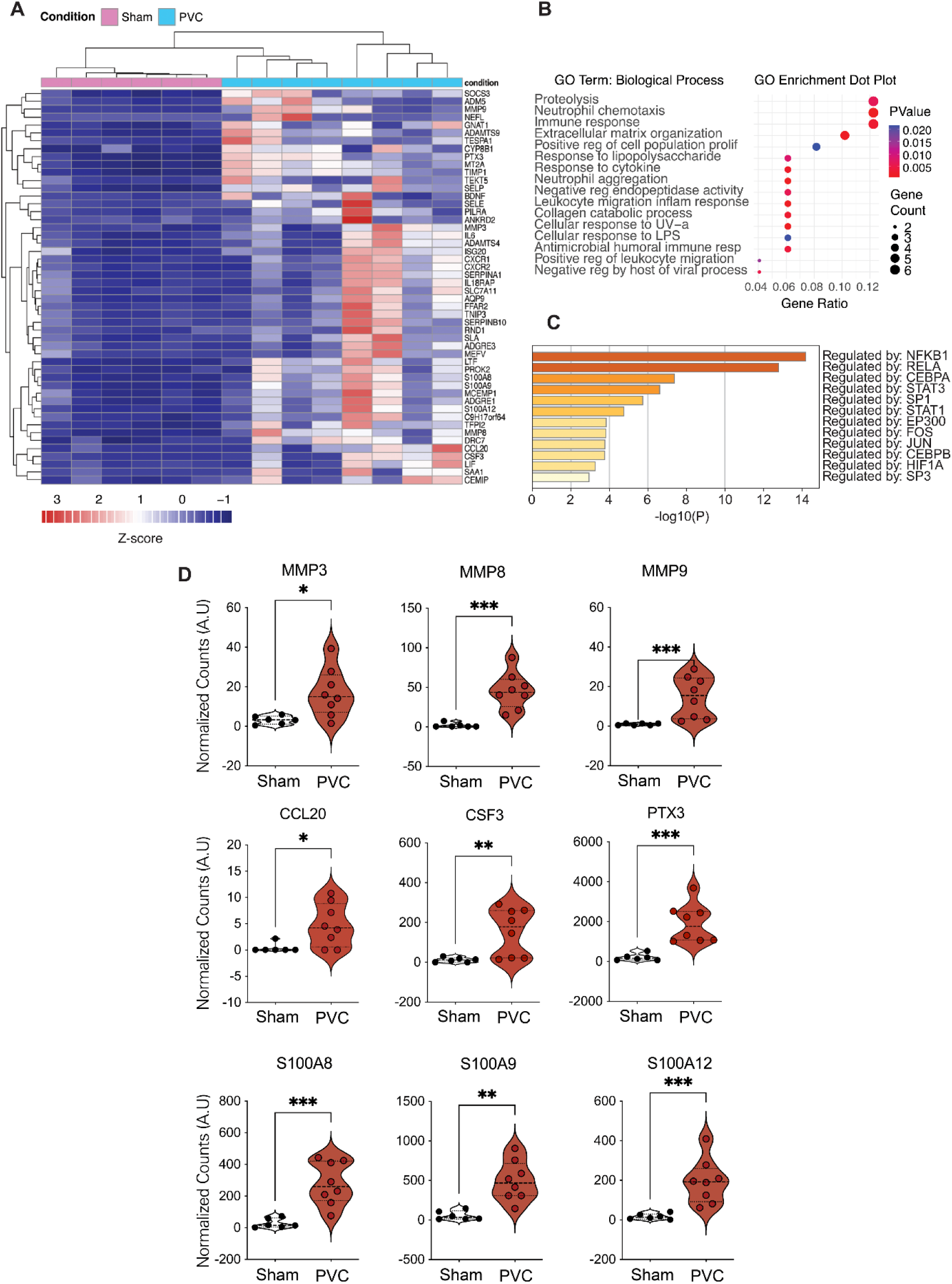
PVC-CM results in a distinct transcription profile. RNAseq was used to identify differentially transcribed genes. (A) Heatmap displaying the top 50 upregulated genes. (B) Gene set enrichment analysis of the top 50 upregulated genes. (C) Enrichment for transcriptional regulatory networks identified from the transcriptional signature. (D) Violin plots showing transcripts of interest differentially expressed between groups. Non-parametric Mann Whitney test with two-tailed was used for comparison between groups; p-values are shown in asterisks * <0.05, **<0.01, *** <0.001.

Among the top five genes with the highest mRNA expression between conditions were metalloproteinases MMP8 (31-fold increase and adjusted p-value 1.67E-9), MMP9 (28- fold increase and adj p-value 3.58E-4), and MMP3 (6-fold increase and adj p-value 0.007) (Fig. 8D). Other metalloproteinases such as ADAMTS9 and ADAMTS4 are also upregulated, as well as the MMP inhibitor TIMP1 (Fig. 8A). This upregulation pattern is consistent with the cardiac remodeling described above in the PVC-CM model. We identified upregulation of numerous genes linked to macrophage activation, immune cell recruitment, and cytokine signaling/production, including CCL20 (C-C motif chemokine ligand 20; 13-fold increase and adj p-value 0.045), CSF3 (Granulocyte colony-stimulating factor 3; 13-fold increase and adj p-value 0.008), and PTX3 (Pentraxin 3; 9-fold increase and adj p-value 2.25E-8) (Fig. 8D). Additionally, SELE (E-selectin), CXCR1, CXCR2 (CXC chemokine receptor 1 and 2), ADGRE1 (Adhesion G Protein-Coupled Receptor E1), IL6 (interleukin 6), SOCS3 (Suppressor of cytokine signaling 3) and CCR1 (C-C chemokine receptor type 1) were all upregulated in PVC-CM vs the sham group (Fig. 8A), suggesting that inflammatory and fibrotic responses work together to sustain cardiac remodeling. An additional set of genes of interest that are strongly upregulated in the PVC-CM group are the S100 Ca^2+^ binding proteins S100A12 (12-fold increase and adj p-value 3.20E-6), S100A8 (10-fold increase and adj p-value 1.21E-4), and S100A9 (9-fold increase and adj p-value 6.89E-5) (Fig. 8D). These genes and their protein products are key biomarkers for cardiac disease, playing a pivotal role in the inflammatory and fibrotic responses of the heart to pathological insults.

## Discussion

### Fibrosis in PVC-CM

Fibroblasts can exist in different states and levels of activation^30, 31^. A commonly used marker to assess fibroblast to myofibroblast differentiation is αSMA. This marker is associated with the expression of stress fibers, promoting migration, and scar compaction^29^. In fact, a high percentage of myofibroblasts are VIM^+^ and αSMA^+^ in models of *replacement fibrosis* such as myocardial infarction^32^. Although αSMA immunostaining was intended to identify myofibroblasts, it revealed patterns consistent with pericytes and smooth muscle of blood vessels (Fig. 2A). In contrast, VIM staining within the interstitial cell population was convincingly selective for fibroblasts in our LV samples (Fig. 2A).

Quantification of the VIM^+^ and αSMA^+^ co-localization signal (Fig. 3, Table 2) showed no increase in PVC-CM compared to the sham group. This result suggests that the myofibroblast population did not increase in PVC-CM. Yet, together with an increase in collagen content, we found a doubling of fibroblasts-to-myocyte ratio (VIM^+^ cells) in PVC- CM (Table 1), suggesting that the number of fibroblasts is elevated in PVC-CM. These observations agree with other reports showing that myofibroblasts (αSMA^+^ fibroblasts) are usually not abundant in non-replacement fibrosis^38^ such as the *reactive fibrosis* observed in PVC-CM. Reactive fibrosis usually responds to mechanical and neurohumoral stress. It affects cardiac contractility and promotes reentrant arrhythmias by changing electrophysiological tissue properties, including conduction velocity and ventricular activation patterns^39, 40^. Both local and regional myocardial fibrosis could have significant clinical implications as they are known predictors of ventricular arrhythmias and SCD^4, 41–46^.

### IL-1β in PVC-CM

IL-1β is a cytokine produced by various cell types, including monocytes, macrophages, cardiac fibroblasts, and endothelial cells, and is considered a key player at the intersection between cardiac fibrosis and sterile inflammation^31, 47–50^. Its expression was elevated in the PVC-CM group (Fig. 5A), and several actions of IL-1β on the cardiac tissue described by others correlate with the phenotype observed in our PVC-CM model. For example, IL-1β has been implicated in cardiac hypertrophy^51, 52^, while we recently described cardiac hypertrophy for the PVC-CM model^8, 18^. Additionally, IL-1β decreases L-type Ca^2+^ current^53^, decreases SERCA expression, and reduces the amplitude of Ca^2+^ transients in ventricular myocytes^54^. These effects of IL-1β on Ca^2+^ handling are consistent with the phenotype previously described in our PVC-CM model^7, 55, 56^. Moreover, this PVC-CM model showed a decrease in I_to_ current and a prolongation in the action potential duration^55^, also described previously as effects of IL-1β^57^. These associations support the new hypothesis that IL-1β is an important mediator of the PVC- CM phenotype. Studies in vivo have shown that deletion of IL-1β signaling decreases inflammatory and fibrotic responses in a model of myocardial infarction^35, 58^, arguing that elevated IL-1β may also be involved in the pro-fibrotic effect observed in PVC-CM, a hypothesis that should be tested in future studies.

### Angiogenesis and Eccentric Hypertrophy

An underlying function of activated fibroblasts is to provide mechanical and humoral support to endothelial cells during angiogenic responses^59^. Fibroblasts influence angiogenesis by secreting several angiogenic factors (e.g., VEGF, FGF, angiopoietins among others) and by modulating the stiffness of the extracellular matrix^60–63^. Our previous work showed elevated levels of VEGF-A^18^ and, as shown here, VEGF-B is similarly elevated in LV PVC-CM (Fig. 6A). These factors have complementary functions in the angiogenic response^64^. VEGF-A is a potent master regulator of angiogenesis mainly secreted by cardiomyocytes in the heart tissue^65^, which is important to initiate formation of immature vessels by vasculogenesis and angiogenic sprouting^66^. In contrast, VEGF-B has a modulatory role in improving angiogenesis by promoting the survival of endothelial cells and multiple other cell types^37^. In addition, increased VEGFs have been associated with cardiac hypertrophy^67, 68^. A protein abundant in endothelial cells but also expressed by cardiomyocytes, eNOS, was also elevated in LV samples of the PVC-CM group (Fig. 6A), further supporting the presence of an angiogenic response in PVC-CM.

We previously reported an increase in LV mass index as measured using two echocardiography methods: the area-length and the linear method. This increase occurs without a change in the relative wall thickness, which indicates eccentric hypertrophy^8^. This finding was later confirmed through microscopic examination, which showed an increase in cross-sectional area of the ventricular myocytes in PVC-CM compared to the sham group^18^. These measurements were performed on the same animal cohorts as those used in this publication. With the addition of more data, the values for myocyte cross-sectional area were updated in Table 1. At a tissue level, cardiomyocyte size in situ increased in both width and length^18^, together with increased in cell capacitance measured in isolated ventricular myocytes^55^, findings that are consistent with eccentric hypertrophy in PVC-CM animals. Moreover, molecular markers of pathological hypertrophy (such as store-operated Ca^2+^ entry, calcineurin activity, phospho-NFAT/total NFAT ratio, β-myosin heavy chain, and skeletal type α-actin) were unchanged, providing evidence of an *adaptive* cardiac remodeling^18^. It is well established that transition from adaptive to decompensated cardiac hypertrophy largely evolves from capillary rarefaction^13^. However, in our PVC-CM model, the density of capillaries remained unchanged compared to the sham group (Table 1). Our capillary density values using the IB4 staining (Fig. 6C and Table 1) are comparable to these reported in cardiac tissue by other groups^69, 70^. This lack of rarefaction in our PVC-CM model is achieved by increasing the capillary-to-myocyte ratio by 19% (Table 1), compensating for the volume “taken” by the hypertrophied myocytes. In addition, pericytes are supportive cells key for capillary function including capillary lumen size regulation^71, 72^. Although the number of pericytes-to-myocyte ratio shows a tendency to increase, the absence of statistical significance in this measure (Table 1) may be attributed to instances where a single pericyte is shared between closely spaced capillaries, as observed in the zoomed panels of Fig. 7B. This phenomenon could suggest the formation of new capillaries in the PVC-CM group. Therefore, the increase in capillaries in PVC-CM is required to support the hypertrophy of cardiomyocytes (Table 1), the expansion of other cells types such as fibroblasts in the interstitial space (Fig. 1B), and the heightened oxygen consumption resulting from PVCs^73^. Additionally, capillaries in the PVC-CM group show a clear increase in their cross-sectional area, similar to what was described previously in a hypertrophy model induced by VEGF-B overexpression^74^ and there is also a significant increase in the number of larger blood vessels. All these findings suggest that: 1) vascular resistance should be reduced in the LV tissue, and blood perfusion should be favored in the PVC- CM group, 2) elevated angiogenesis and larger lumen of capillaries maintain appropriate blood supply to balance the metabolic demand of the hypertrophic phenotype, and 3) adequate perfusion should minimize myocyte loss and the need for replacement fibrosis.

### Global transcriptional changes in PVC-CM

The RNAseq data provided further evidence of extracellular matrix remodeling in PVC- CM by showing an upregulation of MMPs (3, 8, and 9), ADAMTS4 and ADAMTS9. These extracellular proteases can be secreted by fibroblasts, endothelial, and smooth muscle cells and are involved in extracellular matrix adaptations as part of many processes such as development, fibrotic, and angiogenic responses^75–77^. ADAMTS4 and MMP9 are secreted by cardiac fibroblasts^78, 79^ and mediate the release of ECM-stored TGF-β, which is an important pro-fibrotic factor^80^. It was shown that the inhibition of ADAMTS4 prevents cardiac fibrosis in a model of pressure-overload in rats^80^, while ADAMTS9 deficiency leads to heart abnormalities related to insufficient cleavage of versican^77^, a large extracellular matrix proteoglycan involved in cardiac fibrosis^80^. Additionally, pro-fibrotic cytokines such as IL-1β and TGF-β can upregulate MMPs expression and activity in cardiac fibroblasts, promoting extracellular matrix remodeling^81^. TIMP1, also elevated in our PVC-CM model, is an endogenous inhibitor of metalloproteinases, having preference for MMP9^82–84^. The complex interplay between MMPs and TIMPs dynamically controls the extracellular matrix homeostasis, where an increase in TIMPs expression is associated with heart disease^84^.

RNAseq experiments identified several transcripts linked to inflammatory responses in PVC-CM. Notably, transcripts such as CCL20, CSF3, and PTX3 were elevated by more than 9-fold. Cytokines such as TGF-β and IL-1β activate the chemokine cascade, including CCL20, which further enhances the chemotaxis of immune cells and contributes to fibrosis^85–87^ and potentially to revascularization^88^. CSF3 is a key cytokine involved in neutrophil infiltration and promotes a profibrotic phenotype in the heart^89^. PTX3 acts as a regulator of the inflammatory response and tissue remodeling, including the differentiation of monocytes into fibrocytes, and its knockdown has been shown to decrease cardiac fibrosis^90, 91^. Additionally, PTX3 has been studied as a biomarker linked to an increased risk of cardiac events in patients with HF^92^. Several additional genes related to inflammation were upregulated in PVC-CM. SELE, which encodes E-selectin, is crucial for immune cell trafficking, as it facilitates the adhesion and transmigration of leukocytes into tissues^93^. Importantly, SELE is not constitutively expressed but is upregulated on the surface of endothelial cells in response to TNF-α or IL-1β^93^. Other notable genes upregulated were CXCR1 and CXCR2, encoding C-X-C Motif Chemokine Receptors 1 and 2, respectively, both of which play a role in neutrophil recruitment^94^. The alarmins S100A8 and S100A9, abundantly stored in neutrophils as a heterodimer^95^, were also significantly upregulated in PVC-CM tissue (Fig. 8D). Cardiomyocytes also express S100A8/S100A9, which have been implicated in mitochondrial dysfunction and cardiomyocyte impairment in ischemia/reperfusion injury models^96^. These alarmins also are important mediators in the fibroblast-macrophage functional interaction leading to cardiac fibrotic remodeling^97^. S100A12 has been less studied in cardiovascular diseases but together with S100A8/9 are considered as potential biomarkers for cardiac diseases^98,99^. Collectively, the gene transcription profile observed in PVC-CM demonstrated a strong enrichment of the NF-κB transcription factor network (Fig. 8C). This network serves as a pivotal hub modulating immune responses and inflammation^100–103^. Consequently, the RNAseq results strongly indicate that an aseptic inflammatory response is associated with the adverse cardiac remodeling observed in PVC-CM. This finding opens new avenues for future research into the underlying mechanisms of PVC-CM.

## Study limitations

In this study, a high burden of PVC of 50% (bigeminy) used in the experimental group is rarely observed in the clinical setting, however, it ensures that all animals will develop cardiomyopathy as previously published^104^, minimizing canine use in research. Like in humans, frequent PVCs decrease the cardiac output with a tendency to produce a mild cardiomyopathy, and the severity of the ventricular dysfunction is proportional to the PVC burden^5, 105^. Similar to the clinical setting, eliminating PVCs results in the recovery of the ventricular function in a matter of weeks^9^. Unfortunately, histological studies in human samples have not been reported, in part because the PVC-CM diagnosis can only be achieved after the LVEF recovers upon elimination of PVCs. Therefore, using this animal PVC-CM model is an effective alternative to perform these histopathological studies in a controlled manner.

Although this large animal model is one of the best models to study PVC-CM, a significant limitation is the implementation of interventional molecular tools and genetic modifications. Therefore, this is mainly a comparative study, and cause-effect relationships must be further tested in simplified models that might lose translational value.

## Clinical implications

This study demonstrates that cardiac remodeling in PVC-CM is characterized by increased angiogenesis and capillary diameter, supporting the hypertrophic response. Transcriptional changes reveal an underlying aseptic inflammatory response, which together with fibroblast proliferation and activation, leads to interstitial fibrosis. Reactive and interstitial fibrosis has been shown to affect electrophysiological properties that are likely associated with worse outcomes as demonstrated by late gadolinium enhancement and extracellular volume assessed by cardiac MRI^43–45, 106^. Similar to tachycardia-induced cardiomyopathy^107, 108^, interstitial fibrosis in PVC-CM could result in increased risk of ventricular arrhythmias and mortality despite resolution of LV dysfunction after ablation of PVCs. A better understanding of the mechanisms driving inflammation and interstitial fibrosis in PVC-CM, as well as in other cardiomyopathies, could help to identify new therapies to prevent or minimize a persistent LV substrate associated with increased mortality and SCD.

## Acknowledgements

This work was partially funded by National Institutes of Health /NHLBI grant R01 HL139874 (Jose F. Huizar) and R01 AR068431 (Montserrat Samsó), Department of Veterans Affairs grant 5I01BX004861 (Jose F. Huizar), SECTEI-Mexico fellowship SECTEI/135/2019 (J. M. Lourdes Medina-Contreras) and a postdoctoral fellowship from the American Heart Association and the D.C. Women’s Board 836430 (Jaime Balderas-Villalobos). Microscopy was performed at the VCU Department of Anatomy and Neurobiology Microscopy Facility, supported, in part, with funding from the NIH-NINDS Center core grant (5P30NS047463).

## Author Contributions

JM, JB, MS, JFH, JME Conceived and designed research; JM, JB, JH, KK, AT, MS, JG performed experiments; JM, JB, JH, JG analyzed data; JM, JB, JFH, JG, JME interpreted results of experiments; JM, JB, JG, JME prepared figures; JM, JG, JME drafted manuscript; JB, MS, KK, AT, JFH edited and revised manuscript; JM, JB, JH, KK, AT, MS, JG, JFH, JME approved final version of manuscript.

## Data availability statement

The data supporting this study’s findings are available from the corresponding author upon reasonable request.

The authors have no conflicts of interest to declare.

## Abbreviations

PVCs =: premature ventricular contractions.
SCD =: sudden cardiac death
LV =: left ventricle
PVC-CM =: PVC-induced cardiomyopathy
NBF =: neutral buffered formalin
PBS =: phosphate-buffered saline
TEM =: transmission electron microscopy
αSMA =: α-smooth muscle actin
VIM =: vimentin
IB4 =: isolectin B4
IL-1β =: Interleukin 1β
VEGF =: vascular endothelial growth factor
WGA =: wheat germ agglutinin
eNOS =: endothelial nitric oxide synthase

