## supplementary material for "High burden of premature ventricular contractions upregulates transcriptional markers of inflammation and promotes adverse cardiac remodeling linked to cardiomyopathy"

**Supplemental material**

**Methods**

**Protein extraction & Western blotting**

Antibodies used and incubation conditions are described in Table S1

*Alpha smooth muscle actin ($\alpha$SMA)*

LV free wall frozen samples were pulverized in a mortar under liquid nitrogen, followed by collagenase digestion using Worthington type II collagenase (1 mg/ml), then reconstituted in Hank's balanced salt solution (HBSS) with 0.5 mM CaCl2 during 14 min at 37°C with gentle rocking. Samples were then sonicated using myosin extraction buffer (MEB) prepared as follows: 300 NaCl, 100 $NaH_2PO_4$, 50 $Na_2HPO_4$, 20 $Na_4P_2O_7$, 1 MgCl2, 10 EDTA, 0.5 EGTA, 1 DTT, 50 NaF, 1 PMSF, 1 Orthovanadate (in mM, pH 6.5), supplemented with 0.5% NP-40 and protease inhibitors (cOmplete$^{TM}$ Roche). After lysis, homogenates were centrifuged at 8,400 g and the soluble fraction was recovered and stored at -80°C for later use.

*Vimentin, IL-1β and eNOS*

Liquid nitrogen-frozen pulverized LV free wall tissue was homogenized using a traditional RIPA lysis buffer (25 Tris-HCl, 150 NaCl, 2 EDTA, 0.5 EGTA, 1 PMSF, 1 Orthovanadate, 50 NaF [in mM, pH 7.6]) supplemented with 1% NP-40, 0.5% sodium deoxycholate, 0.1% SDS, and protease inhibitors [cOmpleteTM, Roche]), followed by centrifugation at 11,600 g; the soluble fraction was recovered stored at -80°C for later use.

*VEGF-B*

Pulverization of the LV free wall tissue was made as described previously, then mixed and vortexed with sucrose-tris-magnesium (STM) lysis buffer (250 Sucrose, 50 Tris-HCl, 5 $MgCl_2$ [in mM, pH 7.4]) supplemented with protease inhibitors. Then, homogenates were subjected to consecutive centrifugations: 800g, 15'. This supernatant1 was centrifuged at 6,000 g, 15'. Supernatant 2 was ultracentrifuged at 43,000 g for 1h. Supernatant 3 was the cytosolic fraction.

The determination of protein concentration was assessed using the Bradford assay (Sigma) and samples were aliquoted and immediately stored at -80°C until their use. Samples were subjected to SDS- polyacrylamide gel electrophoresis and transferred to PVDF membranes. The blots were blocked with 5% non-fat milk (BioRad) in 0.1% TBS-Tween and incubated overnight at 4°C with the primary antibodies indicated in Table S1. After a few washes and secondary antibody incubation, blots were exposed to an enhanced chemiluminescent (ECL) assay (Invitrogen) and images were obtained in the FluorChem R (ProteinSimple, CA) gel imager. GAPDH was used as loading control for each blot. Images were analyzed using the software AlphaView (ProteinSimple, CA).

**Immunofluorescence**

Antibodies used and incubation conditions are described in Table S2

Tissue slices (40 µm thick; Leica vibratome V1 1000S) obtained from 10% neutral buffered formalin (NBF)-fixed LV free wall were subjected to antigen retrieval at pH 9 (abcam, ab93684), then incubated with primary antibodies or lectins described in Table

S1. This was followed by appropriate secondary antibody and DAPI incubation, washed with PBS, placed in slides, and mounted using ProLong Gold (Invitrogen) media. Cardiac tissue was examined using confocal laser scanning microscopy (Zeiss LSM 700). Colocalization analysis is described in the supplementary material.

**Collagen determinations using Sirius Red Fast Green staining in situ.**

The 10% NBF fixed tissue was paraffin embedded, then sectioned into 10 µm-thick slices (Reichert-Jung 820 II, rotary microtome), rehydrated, and stained with Sirius Red /Fast Green to assess collagen fiber deposits[1]. Micrographs were acquired at 20X using a Carl Zeiss microscope (Axio Imager Z2) equipped with a color camera. Briefly, the images were converted to an RGB stack, and the red channel was used to detect collagen fibers stained with Sirius Red. The red-stained collagen fibers were selected by using the threshold function of ImageJ software and interstitial fibrosis was reported as percentage of the area covered by collagen fibers in the whole image.

**Artificial intelligence powered analysis.**

The artificial intelligence-powered autonomous cell analysis software, AIVIA Go (Leica Microsystems, Bellevue, WA), was used to perform morphometric analysis of images by identifying and selecting patterns compatible with interstitial cells, capillaries, blood vessels, and cardiomyocytes. The software, which uses a machine learning-powered *pixel classifier* tool, was trained to recognize unique features of each structure or cell type according to size, shape, and signal intensity. Once *regions of interest* were established, the pixel classifier was automatically applied to all images in a batch mode. This process generated image segmentation and ensured a consistent identification process. Finally, a script was created using the software console to perform automatic quantification of parameters such as quantity and dimensions (surface area) of the *regions of interest* in each image.

**Colocalization analysis**

Confocal images co-stained using ant-$\alpha$SMA and anti-vimentin antibodies were subjected to colocalization analysis. The Pearson and Mander coefficients along with the percentage of overlapping pixels for $\alpha$SMA and vimentin channels were estimated using the *Colocalization Finder* plug-in on ImageJ software. Briefly, after splitting the channels, the histogram of the images was adjusted to maximize the dynamic range and the colocalization thresholds were set for both channels to exclude pixels with intensity at background level. The colocalization parameters were measured for several images per animal and differences between groups were studied using hierarchical analysis (nested t-test).

**Transmission Electron Microscopy.**

Samples were processed and stained using 2% osmium tetroxide and saturated uranyl acetate as previously described[2]. Images were acquired on a Tecnai F20 or a Jeol JEM-1230 transmission electron microscope, both equipped with a Gatan UltraScan 4K x 4K CCD camera.

Table S1
Western blot:

| Antibody | Protein load | SDS-PAGE | Antibody dilution | Catalog number |
|---|---|---|---|---|
| anti-αSMA | 50 μg/well | 8% | 1:1000 | MA5-41117, Invitrogen |
| anti-Vimentin | 20 μg/well | 8% | 1:500 | MA5-11883, Invitrogen |
| anti-VEGF B | 30 μg/well | 15% | 1:1000 | 2463, Cell Signaling |
| anti-eNOS | 60 μg/well | 8% | 1:1000 | PA1-037, Invitrogen |

Table S2
Immunofluorescence:

| Antibody / marker | Dilution / Concentration | Incubation time | Catalog number |
|---|---|---|---|
| anti-αSMA | 1:200 | 4°C, over night | MA5-41117, Invitrogen |
| anti-Vimentin | 1:50 | 4°C, over night | MA5-11883, Invitrogen |
| IB4-AF488 | 20 μg/ml | 4°C, over night | I21411, Invitrogen |
| WGA-AF633 | 14 μg/ml | 4°C, over night | Ab76003, Abcam |
| anti-Mouse-AF488 | 1:100 | RT, 3 h | A-11001, Invitrogen |
| anti-Rabbit-AF647 | 1:200 | RT, 3 h | A-21244, Invitrogen |
| anti-Mouse-AF568 | 1:100 | RT, 3 h | A-11031, Invitrogen |

*RT, room temperature
